# THAP1 Modulates Oligodendrocyte Maturation by Regulating ECM Degradation in Lysosomes

**DOI:** 10.1101/2020.09.27.316133

**Authors:** Dhananjay Yellajoshyula, Samuel S. Pappas, Abigail Rogers, Biswa Choudhury, Xylena Reed, Jinhui Ding, Mark R. Cookson, Vikram Shakkottai, Roman Giger, William T. Dauer

## Abstract

Mechanisms controlling myelination during CNS maturation play a pivotal role in the development and refinement of CNS circuits. The transcription factor THAP1 is essential for timing the inception of myelination during CNS maturation through a cell-autonomous role in the oligodendrocyte lineage. Here, we demonstrate that THAP1 modulates ECM composition by regulating glycosaminoglycan (GAG) catabolism within oligodendrocyte progenitor cells (OPCs). *Thap1^-/-^* OPCs accumulate and secrete excess GAGs, inhibiting their maturation through an auto-inhibitory mechanism. THAP1 controls GAG metabolism by binding to and regulating the *GusB* gene encoding β-glucuronidase, a GAG-catabolic lysosomal enzyme. Applying GAG-degrading enzymes or overexpressing β-glucuronidase rescues *Thap1^-/-^* OL maturation deficits *in vitro* and *in vivo.* Our studies establish lysosomal GAG catabolism within OPCs as a critical mechanism regulating oligodendrocyte development.

## INTRODUCTION

The oligodendrocyte (OL) lineage plays a critical role in CNS development and function (Almeida and Lyons, 2017; Bergles and Richardson, 2015). The OL lineage in its mature cell state is responsible for wrapping axons with an insulating myelin sheath that supports rapid neurotransmission (Waxman and Bennett, 1972). Developmental myelination during a critical period of post-natal development plays an important role in the establishment of neural circuits (Fields, 2008; Rice and Barone, 2000; Wang et al., 2018). Regulated myelination in the adult CNS is also increasingly recognized to play an essential role in circuit plasticity, including motor learning (Bacmeister et al., 2020; Bergles and Richardson, 2015; McKenzie et al., 2014; Pepper et al., 2018; Xiao et al., 2016). Mechanisms regulating OL maturation are therefore central to the development of neuronal circuits as well as their plasticity and maintenance.

OL differentiation leading to myelination is regulated by cell-intrinsic pathways (Elbaz and Popko, 2019) and extrinsic signals that emanate from axons and the surrounding extracellular matrix (ECM) (Colognato and Tzvetanova, 2011; Harlow and Macklin, 2014; Sandvig et al., 2004). The ECM is a complex three dimensional structure surrounding cells, providing both a physical scaffold and a growth factor organizing center orchestrating signaling cascades that regulate mobility, development and function (Colognato and Tzvetanova, 2011; Fawcett et al., 2019; Theocharidis et al., 2014). The ECM is composed of glycosaminoglycans (GAGs), a class of long unbranched mucopolysaccharides, GAG-modified proteoglycans (GAG-PG) and fibrous proteins (e.g., collagen, elastin) (Esko et al., 2009). Considerable literature demonstrates that GAGs in their free form and as proteoglycans (e.g., chondroitin sulfate proteoglycan; CSPG) inhibit OL maturation both *in vitro* and *in vivo,* demonstrating that GAGs powerfully regulate OL development (Karus et al., 2016; Keough et al., 2016; Kuboyama et al., 2017; Lau et al., 2012; Pu et al., 2018; Siebert and Osterhout, 2011; Sloane et al., 2010).

GAG and proteoglycan content is determined through a balance of synthesis, secretion, and catabolism (Esko et al., 2009; Nowicka and Greda, 2019; Song and Dityatev, 2018). In addition to the extracellular degradation of ECM components by secreted proteases (Berezin et al., 2014), receptor and cell surface proteoglycans and GAGs are endocytosed and enzymatically catabolized in lysosomes (Rainero, 2016; Rappaport et al., 2016; Winchester, 2005). Failure of intracellular GAG catabolism results in mucopolysaccharidoses, a class of lysosomal storage disorders (Fiorenza et al., 2018; Kobayashi, 2019; Winchester, 2005). Despite their importance, the sources and cellular mechanisms regulating GAG content and composition in the CNS are ill defined. Most studies on the cellular origins of ECM have focused on CSPG synthesis from astrocytes (Jones et al., 2003; Sandvig et al., 2004; Song and Dityatev, 2018). Study of ECM metabolism within the OL lineage itself has been limited (Courel et al., 1998; Garwood et al., 2004; Song and Dityatev, 2018; Yim et al., 1993), despite the likely importance of these processes for OL biology in normal development and disease states.

We demonstrated a cell autonomous role for the transcription factor THAP1 in regulating the maturation of OPC to mature myelinating OLs (Yellajoshyula et al., 2017). Loss-of-function mutations in the *THAP1* gene cause the neurodevelopmental disorder DYT6 dystonia, directly implicating these events in dystonia pathogenesis (Fuchs et al., 2009). Conditional deletion of *Thap1* from the OL lineage significantly impairs CNS myelination by retarding the transition of OLs to mature myelin producing cells, without overtly altering the number of oligodendrocyte progenitor cells (OPC)(Yellajoshyula et al., 2017). The marked delay in myelination corresponds to the development of motor dysfunction in the *Thap1* mutants. The mechanism whereby THAP1 regulates OL lineage maturation was not defined in this study.

Here, we identify a direct role for THAP1 in regulating GAG catabolism and demonstrate directly that dysregulation of GAG catabolism within OPCs impairs their maturation. *Thap1^-/-^* OPCs auto-inhibit their own maturation by accumulating and secreting excess GAGs. LC/MS analyses of OPC conditioned media demonstrate they predominantly secrete chondroitin 4-sulfate, which is significantly increased in *Thap1^-/-^* cells. THAP1 regulates GAG catabolism by binding to and regulating the *GusB (MPS7)* gene encoding the lysosomal GAG-catabolizing enzyme β-glucuronidase. Treatment with GAG degrading enzymes or overexpressing β-glucuronidase rescues the *Thap1^-/-^* OPC deficits in OL maturation and CNS myelination, establishing that excess GAGs mediate maturation delay in the *Thap1^-/-^* oligodendroglial lineage. These findings newly establish GAG catabolism as a major cell intrinsic regulatory pathway driving OL maturation.

## RESULTS

### THAP1 regulates ECM pathway composition during OL development

We demonstrated previously that THAP1 plays a cell-autonomous role in regulating myelination during early postnatal CNS development (Yellajoshyula et al., 2017). To begin to examine the mechanisms responsible for this role, we performed RNAseq at multiple time points on cultured control *(Thap1^+/+^)* and *Thap1^-/-^* (derived from *Thap1*^flx/-^; Olig2-*Cre^+^*) OPC under differentiating conditions. Significantly fewer *Thap1^-/-^* OPC matured to express myelin basic protein (MBP+) following 4 days *in vitro* (DIV4) of treatment with 40 μg/ml T3 thyroid hormone (Fig. 1A-B; Control *(Thap1^+/+^)* = 79.44% ± 4.58; *Thap1^-/-^* = 83.28% ± 6.83; t-test p < 0.001). MBP+ *Thap1^-/-^* OLs are also significantly smaller (Fig. 1A,D) and exhibit less intense MBP staining than controls (Fig. 1A,C). We isolated total mRNA from progenitor (OPC) and differentiating OLs (DIV2 & 4) from three clonal lines each for *Thap1^+/+^* and *Thap1^-/-^* genotypes (Fig. 1E). We used DESeq2 for the RNAseq data derived from 18 samples to identify THAP1-dependent differentially expressed genes (DEG), requiring a greater than log2 1.5 fold change and p < 0.05 significance by the Benjamini-Hochberg false discovery procedure. These analyses identified 1132 (OPC), 703 (DIV2), and 1059 (DIV4) DEGs (Fig. 1G; Table S1).

**Figure 1.**
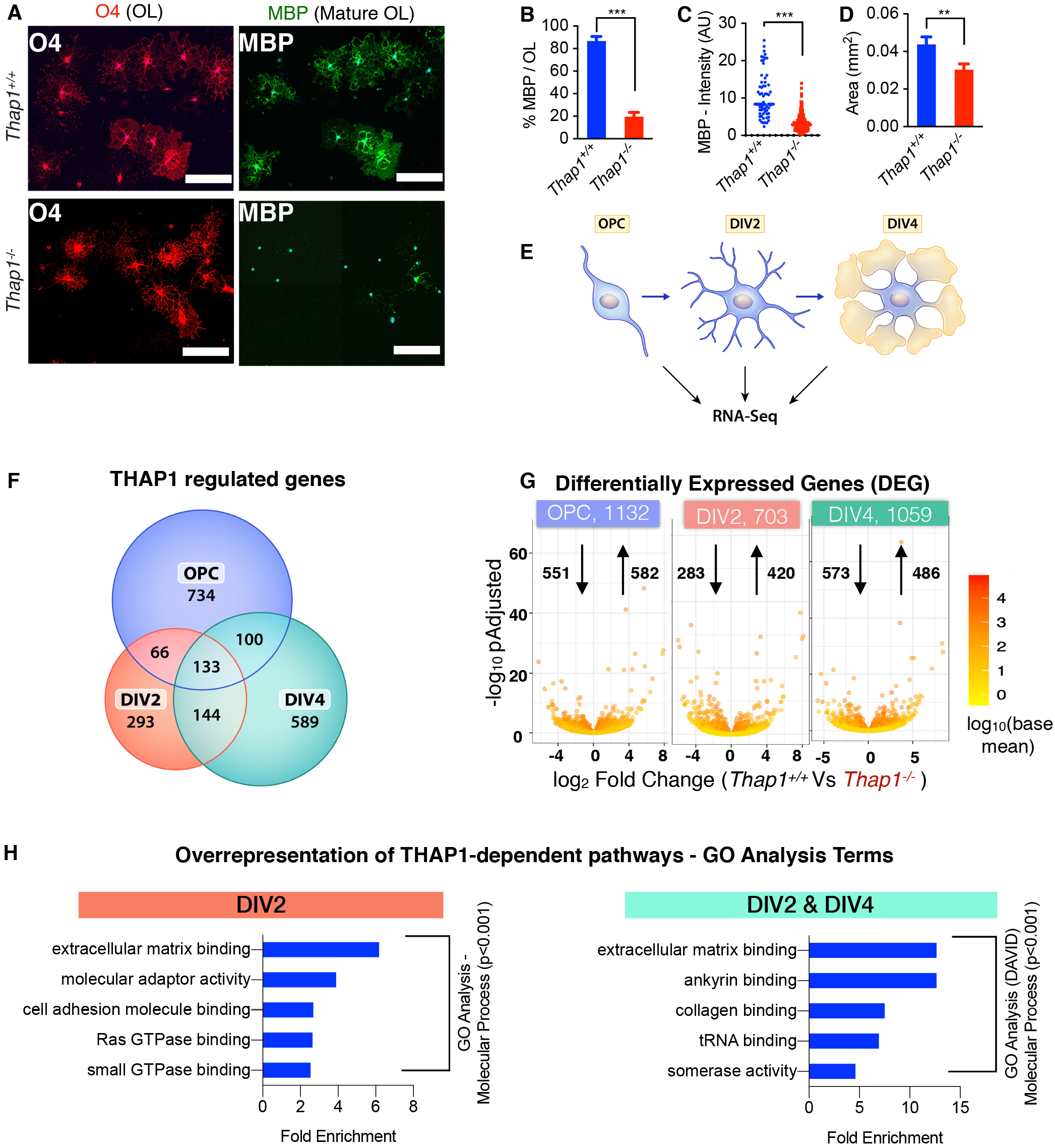
THAP1 loss disrupts the extracellular matrix transcriptome. (A-D) Loss of THAP1 causes maturation deficits in differentiating OPC. (A) Representative Images of control *(Thap1^+/+^)* and THAP1 null (derived from *Thap1*^flx/-^; Olig2-*Cre^+^*) OL differentiated (+T3) for 4 days (scale bar 100 μm). Cultures are stained for O4, MBP, & DAPI. (B) Quantification of the percentage of O4+ cells expressing MBP. Bar graph shows mean values ± SEM. *Thap1^+/+^* = 86.78% ± 3.98 ; *Thap1^-/-^* = 19.58% ± 3.88; t-test: t_(4)_ = 12.07; p = 0.0003 ; N = 3 clonal lines (100 cells per clone) per genotype. (C) Mean intensity of MBP staining in O4+ cells represented as arbitrary units (AU), normalized to background. Each data point in the scatter plot represents MBP+ staining intensity for individual cells. *Thap1^+/+^* = 10.76 AU ± 0.76; *Thap1^-/-^* = 3.32 AU ± 0.108; t-test: t_(427)_ = 18.21; p < 0.0001. (D) Average area of O4+ cells. Bar graph shows mean values ± SEM. *Thap1^+/+^* = 0.043 mm^2^ ± 0.00402; *Thap1^-/-^=* 0.030 mm2 ± 0.00309; t-test: t_(633)_ = 2.639; p = 0.008. (E) Schematic illustrating RNAseq analyses. Total RNA was extracted from three independent clonal lines each of *Thap1^+^*^/+^ and *Thap1^-/-^* cells at three different time points (OPC; DIV2 and DIV4) and analyzed using Illumina Hiseq (Methods). (F) Venn diagram depicting the overlap of *Thap1^+/+^* vs *Thap1^-/-^* differentially expressed genes (DEG) from OPC, DIV2 & DIV4. (G) Number of differentially expressed genes (DEG) at each time point analyzed. X-axis = log2 fold change (magnitude of change for the DEG) in *Thap1^-/-^.* Y-axis = -log10 pAdjusted depicts the statistical significance of the change. The number of up- and down-regulated genes are indicated by arrows. (H) The DEG at either DIV2 (703) or both DIV2 & DIV4 (277) were used for enrichment analysis of GO (Gene Ontology) terms to identify overrepresented biological pathways using both GENEONTOLOGY (http://geneontology.org) and DAVID (https://david.ncifcrf.gov/home.jsp). Graphs shows the most significantly overrepresented GO terms (p<0.001, sorted as fold overrepresented) in OL from DIV2 (GENEONTOLOGY, GO - Molecular Function), and from the DIV2 & DIV4 overlapping group (DAVID, GO - Molecular Function).

Analyses of the RNAseq data using estimated sample distance (Heat map, Fig. S1A) and principal component analysis (Fig. S1B) demonstrated that, within the *same* genotype, differentiating OL (DIV2 & 4) were distinctly different from OPC. Thus we focused on DIV2 & 4 cells, identifying two groups of THAP1-dependent differentially expressed genes (DEG): those in immature (DIV2) cells, and those persisting through maturation (i.e., differentially expressed at both DIV2 & 4) (Fig. 1F). There were 703 DEG limited to DIV2 and 277 DEG present at both DIV2 & 4 (Fig. 1F). These DEG were used for enrichment analysis of GO (Gene Ontology) terms to identify overrepresented biological pathways using both GENEONTOLOGY (http://geneontology.org) and DAVID (https://david.ncifcrf.gov/home.jsp) (Table S2). GO enrichment analyses (see Methods) demonstrated significant overrepresentation of ECM-related pathways in both groups of genes (Fig. 1H, S1C & Table S2). Specific pathways identified were (1) GAG metabolism: (2) ECM-binding (including members of integrin and collagen families): (3) signaling components (Rho GTPases): and (4) cytoskeletal polymerization & filopodium assembly (Fig. 1H & S1C). These analyses suggested strongly that ECM pathways are dysregulated in the *Thap1^-/-^* oligodendroglial lineage.

### Accumulation of Glycosaminoglycan in *Thap1^-/-^* oligodendrocytes

Guided by the strong representation of ECM and particularly GAG metabolism pathways, we explored whether *Thap1^-/-^* cultures exhibited abnormalities in the levels of GAGs. We focused on two species most closely linked to oligodendrocyte biology: chondroitin sulfate (CS) and hyaluronate (HA-GAG) (Karus et al., 2016; Siebert and Osterhout, 2011; Sloane et al., 2010). Immunostaining of non-permeabilized mature (DIV4) cultures demonstrated that *Thap1^-/-^* cells exhibited significantly increased staining for CS- and HA-GAG (Fig. 2A-D) and prominent accumulation around smaller OL processes (Fig. S2A). Staining specificity was confirmed by treating the cultures with chondroitinase ABC (ChABC) or hyaluronidase, which removed the relevant signals (Fig. 2 A-D). *Thap1^-/-^* cultures contained a greater number of positive cells as well as significantly greater intensity of staining per cell (Fig. 2 A-D). Measurement of bulk GAG content using the carbazole assay confirmed significantly increased GAGs in *Thap1^-/-^* cultures (Fig. S2B; OPC; *Thap1^+/+^* = 0.066 AU ± 0.0006; *Thap1^-/-^* = 0.094 AU ± 0.0022; t-test: t_(2)_ = 12.70; p = 0.006 & OL - *Thap1^+/+^* = 0.036 AU ± 0.0023; *Thap1^-/-^* = 0.059 AU ± 0.0019; t-test: t_(2)_ = 7.96; p = 0.015). These findings are consistent with the RNA-Seq data indicating that THAP1 participates in a GAG metabolism-related pathway.

**Figure 2.**
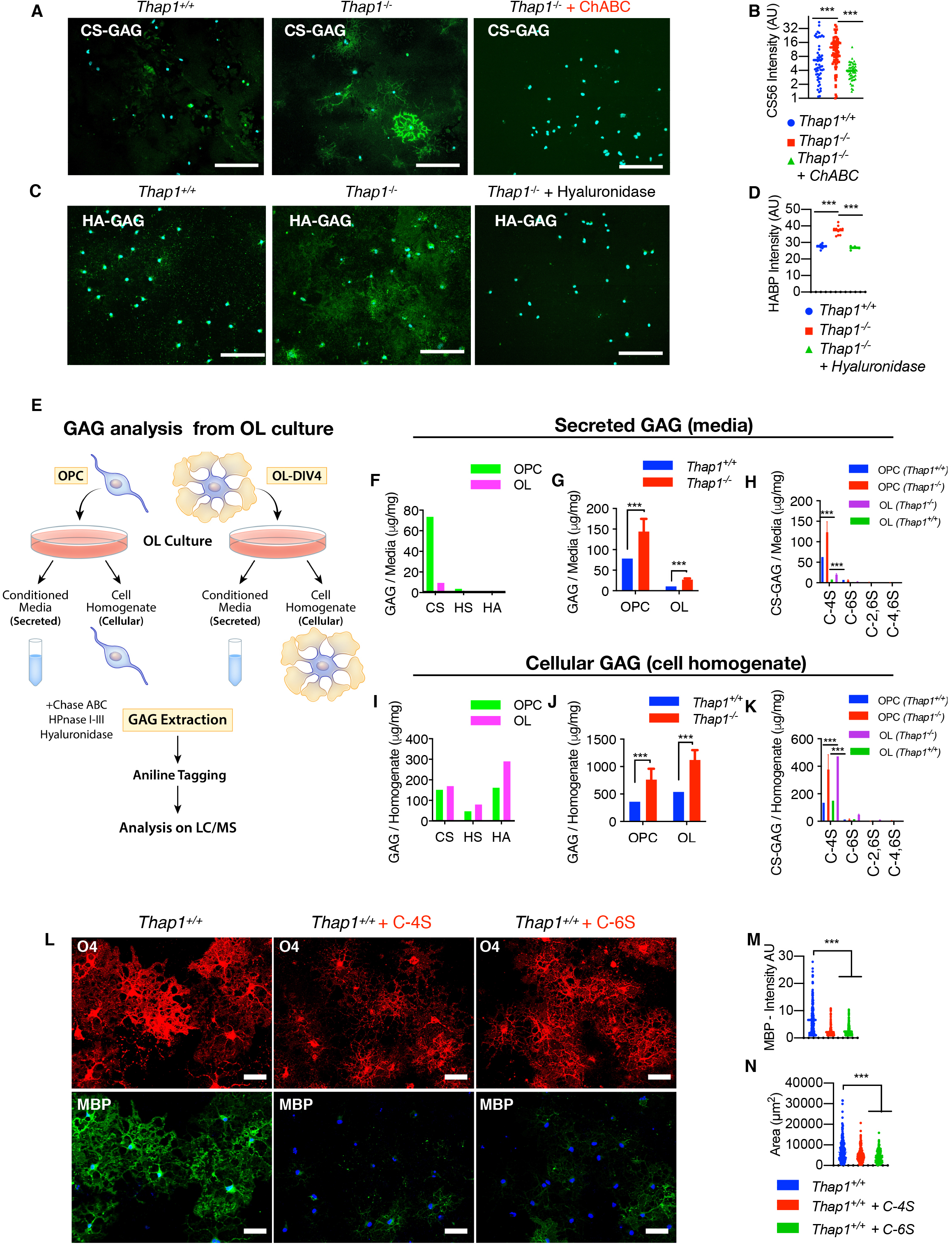
*Thap1* null oligodendrocytes accumulate and secrete excess glycosaminoglycans. Representative images and corresponding quantification for *Thap1^+/+^* and *Thap1^-/-^* OL cells (DIV4) immunostained with (A, B) CS-56 (CS-GAG), and (C, D) biotinylated HABP (HA-GAG) under non-permeabilizing conditions. *Thap1^-/-^* OL exhibit significant accumulation of CS-GAG *(Thap1^+/+^* = 6.68 ± 1.13; *Thap1^-/-^* = 12.60 ± 1.29; *Thap1^-/-^* + ChABC = 3.93 ± 0.286; One-way ANOVA F_(2,205)_ = 14.59, p<0.0001, Dunnett’s multiple comparisons test: adjusted p value < 0.0001) and HA-GAG *(Thap1^+/+^* = 27.64 AU ± 0.38; *Thap1^-/-^* = 37.45 AU ± 2.68; *Thap1^-/-^* + Hyaluronidase = 26.77 AU ± 1.16; One-way ANOVA F_(2,21)_ = 75.59, p<0.0001, Dunnett’s multiple comparisons test: adjusted p value = 0.0004 & < 0.0001). (E) Schematic illustrating the study design for mass spectrometry analyses of CS, HS and HA GAGs using GRIL LC/MS analysis. Total CS, HS and HA GAGs (μg/mg) (F) secreted in the media or (I) from cell homogenate from *Thap1^+^*^/+^ OPC and OL (see Fig. S2C-D for GAG content). (G-H, J-K) Abnormal accumulation of GAGs in *Thap1^-/-^* cultures. Total GAGs (μg/mg) (CS, HA and HA-GAGs combined) (G) secreted in the media or (J) from cell homogenate from *Thap1^+/+^* and *Thap1^-/-^* OPC and OL (see Fig. S2C-D for GAG content). CS-GAGs (μg/mg) and the composition of mono (C-4S, C-6S) and bi-sulfated (C-2,6S and C-4,6S) (H) CS-GAGs secreted in the media or (K) from cell homogenate from *Thap1^+/+^* and *Thap1^-/-^* OPC and OL (see Fig. S2E-F for GAG content). (L-N) C-4S and C-S6S inhibit OL differentiation. Representative Images (L) and corresponding quantification (M-N) of differentiated wild type OPC without (L, left column) or with C-4S (10μg/ml; middle column) or C-6S (10μg/ ml; right column) GAG applied to the culture media from DIV2 to DIV5 (scale bar 50 μm). (M) Mean intensity of MBP staining at DIV5 normalized to background; *Thap1^+/+^* = 6.64 AU ± 0.36 ; *Thap1^+/+^* + C-4S= 2.11 AU ± 0.126; *Thap1^+/+^* + C-6S= 2.33 AU ± 0.128; One-way ANOVA F_(2,681)_ = 119.9, p<0.0001, Dunnett’s multiple comparisons test: adjusted p value = 0.0004 & < 0.0001. Each data point in the scatter plot represents MBP+ staining intensity for individual cells. (N) Area of O4 stained OL *(Thap1^+/+^* = 8038 μm^2^ ± 463.8 ; *Thap1^+/+^* + C-4S= 5122 μm^2^ ± 230.3; *Thap1^+/+^* + C-6S= 4383 μm^2^ ± 198.6; One-way ANOVA F_(2,627)_ = 34.1, p<0.0001, Dunnett’s multiple comparisons test: adjusted p value = < 0.0001). Each data point in the scatter plot represents the area measured for individual cells.

### Chondroitin-4-sulfate is the major GAG elevated in *Thap1^-/-^* OL

Little is known regarding the contribution of the OL lineage to the ECM, such as its ability to produce and secrete GAGs or the identity of those produced. Several species of GAGs (CS, HS and HA) regulate CNS development, including OL lineage maturation (Harlow and Macklin, 2014; Karus et al., 2016; Properzi et al., 2008; Siebert and Osterhout, 2011; Sloane et al., 2010). To determine if GAG species accumulate within THAP1 mutant OL and to characterize their identity, we employed glycan reductive isotope labeling (GRIL) combined with liquid chromatography tandem mass spectrometry (LC/MS) to quantify CS, HS and HA GAGs (Lawrence et al., 2008) in cell homogenates (intracellular) and supernatant (secreted) from DIV4 *Thap1^+/+^* and *Thap1^-/-^* cultures (schematic, Fig. 2E). These analyses demonstrated that there are striking changes in the levels of secreted GAGs during OL maturation (Fig. 2F & S2C). Media from *Thap1^+/+^* OPC cultures exhibited ~8-fold higher GAG levels than in mature OLs, over 95% of which is CS-GAG (Fig. 2F & S2C). In contrast, while significant differences in the composition of GAG were observed in cell homogenates (Fig. 2I & S2D), the relative differences were far less than those observed in the media. THAP1 depletion significantly increased both secreted and cellular GAG (CS, HS and HA combined) levels (Fig. 2G,J & S2C-D). These differences were primarily driven by increases in CS-GAG, the dominant GAG species observed (Fig. 2F & S2C). Levels of HA- & HS-GAGs were also significantly increased in *Thap1^-/-^* cultures (Fig. S2C-D).

The CS-GAG family includes many species with differing amounts and patterns of sulfation that can be detected and resolved by LC/MS. In *Thap1^+/+^* cultures, we found that the dominant form of CS-GAG secreted by OL cultures is chondroitin-4-sulfate (“C-4S”), constituting ~85 - 90% of CS-GAG secreted by OL cultures (Fig. 2H,K & S3E-F). Chondroitin-6-sulfate (“C-6S”) was the only other abundant species (~10% of CS-GAGs) with negligible amounts of non- and bi-sulfated species (“C-2,6S” & “C-4,6S”) (Fig. 2H,K & S3E-F). Consistent with the immunostaining data (Figs. 2A-B), C-4S and C-6S were significantly increased in *Thap1^-/-^* cultures (Fig. 2H,K & S3E-F). We directly tested the ability of these forms of CS-GAG to alter oligodendroglial lineage progression by applying these species to differentiating (wild type) OPC cultures (Figs. 2L-N). C-4S and C-6S (10 μg/ml) each caused a dose dependent inhibition of OL maturation, impairing the progression to mature MBP+ OLs and causing mature cells to occupy a smaller area (Fig. 2L-N). These data demonstrate that the specific species of CS-GAG elevated in *Thap1^-/-^* cultures impair oligodendroglial maturation.

### Accumulation of secreted CS-GAG contributes to *Thap1^-/-^* maturation deficits and exerts an inhibitory paracrine effect

To further explore the hypothesis that increased CS-GAG secretion is responsible for the maturation impairments of *Thap1^-/-^* cultures, we tested the ability of the CS-GAG degrading enzyme ChABC to rescue this phenotype. Consistent with our hypothesis, treatment of *Thap1^-/-^* cultures with 0.1U/ml ChABC significantly increased the proportion of myelinating OL, by increasing the number MBP+ cells (~3-fold increase), the intensity of MBP staining (~3-fold increase), and OL size (~2-fold increase in cell surface area) (Fig. 3A-D). Since *Thap1^-/-^* OLs secrete excess CS-GAGs (Fig. 2), we hypothesized that they may exert non-cell autonomous paracrine effects on neighboring cells. We tested this idea by exploring the ability of *Thap1^+/+^* OPC (GFP-labeled with lentivirus; “LV-GFP”) to mature when co-cultured with unlabeled *Thap1^-/-^* or *Thap1^+/+^* OL (Figs. 3E & S3; cultures contain 1 *Thap1^+/+^*;LV-GFP cell:~10 *Thap1^-/-^* or *Thap1^+/+^* OL). *Thap1^-/-^* OLs significantly impaired the maturation of *Thap1^+/+^*;LV-GFP cells (Fig. 3 F-H). Compared to being cultured in the presence of *Thap1^+/+^* OLs, *Thap1^+/+^* ;LV-GFP OPCs cultured in the presence of *Thap1^-/-^* OLs exhibited significantly lower MBP expression (Fig. 3 F-G; ~ 4-fold lower MBP intensity) and covered a significantly smaller area (Fig. 3 F,H; ~30% smaller area). This inhibitory paracrine effect was eliminated by the addition of ChABC to the media (Fig. 3 F-H). Considered together, these data are consistent with a model whereby *Thap1^-/-^* OLs inhibit the maturation of surrounding cells through a CS-GAG inhibitory paracrine signal.

**Figure 3.**
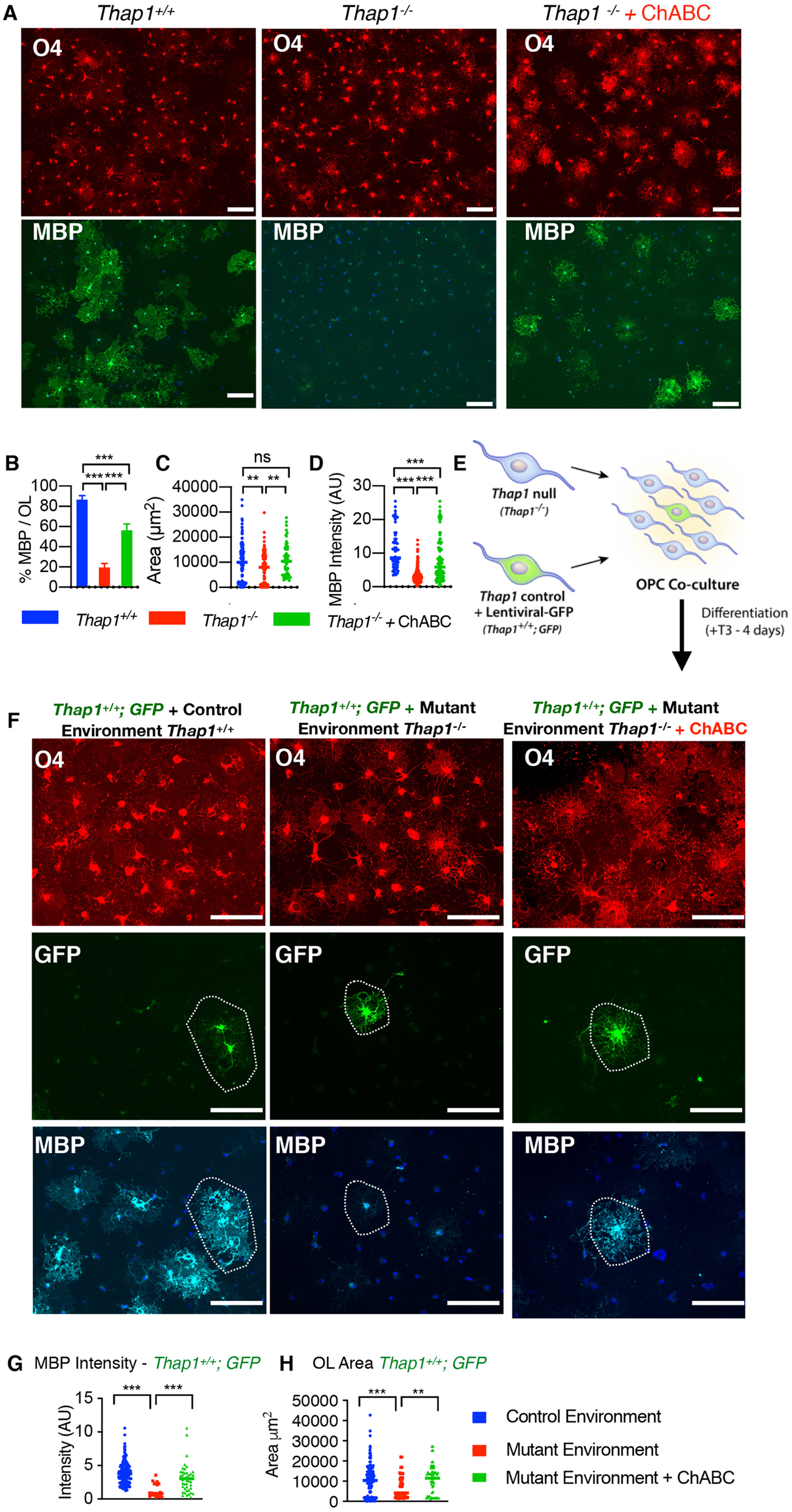
Excess glycosaminoglycan secretion by *Thap1* null cells impairs oligodendroglial maturation via a non-cell autonomous mechanism. (A-D) Treatment with ChABC rescues maturation defects in *Thap1^-/-^* OL. (A) Representative Images (scale bar 100 μm) of *Thap1^+/+^* and *Thap1^-/-^* OL differentiated for 4 days (+T3) either untreated or with ChABC from DIV2 to DIV4 (0.1 U/ml). Cultures are stained for O4, MBP, & DAPI. (B) Quantification of the percentage of O4+ cells expressing MBP. Bar graph shows mean values ± SEM. *Thap1^+/+^* = 85.31% ± 4.14 ; *Thap1^-/-^* = 20.81% ± 4.76; *Thap1^-/-^* + ChABC = 56.29% ± 6.31. One-way ANOVA F<2,27> = 47.94, p < 0.0001, Dunnett’s multiple comparisons test: adjusted p value < 0.0001. (C) Average area of O4+ cells. Each data point in the scatter plot represents the area measured for individual cells. *Thap1^+/+^* = 10163 μm^2^ ± 675.9 ; *Thap1^-/-^* = 7667 μm^2^ ± 549.5; *Thap1^-/-^* + ChABC = 11401 μm^2^ ± 968.3. One-way ANOVA F_(2,309)_ = 6.35, p = 0.002, Dunnett’s multiple comparisons test: adjusted p value = 0.009. (D) Mean intensity of MBP staining in O4+ cells represented as arbitrary units (AU) normalized to background. Each data point in the scatter plot represents MBP+ staining intensity for individual cells. *Thap1^+/+^* = 11.07 AU ± 0.772 ; *Thap1^-/-^* = 3.22 AU ± 0.124; *Thap1^-/-^* + ChABC = 7.86 AU ± 0.63. One-way ANOVA F_(2,524)_ = 134.7, p < 0.0001, Dunnett’s multiple comparisons test: adjusted p value < 0.0001. (E) Schematic illustrating coculture experimental paradigm to test for paracrine effects of *Thap1* null *(Thap1^-/-^)* OL co-cultured with LV-GFP labelled control (*Thap1^+/+^; GFP*) OPC at 10:1 ratio in differentiation media for four days (+T3). (F) Representative images (scale bar 100 μm) of *Thap1^+/+^;GFP OL* cultured in control or mutant environments for 4 days (+T3). Cultures are stained for O4, MBP, & DAPI. (G) MBP+ staining intensity of *Thap1^+/+^; GFP* OL. Each data point in the scatter plot represents MBP+ staining intensity for individual cells. *Thap1^+/+^; GFP* + Control environment = 3.90 AU ± 0.146 ; *Thap1^+/+^; GFP* + Mutant environment = 1.286 AU ± 0.227; *Thap1^+/+^; GFP* + Mutant environment + ChABC = 3.05 AU ± 0.348; One-way ANOVA F_(2,197)_ = 20.26, p < 0.0001, Dunnett’s multiple comparisons test: adjusted p value < 0.0001. (H) Area of *Thap1^+/+^; GFP* OL. Each data point in the scatter plot represents the area measured for individual cells. *Thap1^-/-^* + Control environment = 10163 μm^2^ ± 675.9 ; *Thap1^-/-^* + Mutant environment = 7146 μm^2^ ± 1220; *Thap1^-/-^* + Mutant environment + ChABC = 9151 μm^2^ ± 1160; One-way ANOVA F_(2,266)_ = 6.84, p = 0.0013, Dunnetts’s multiple comparisons test: adjusted p value < 0.0049.

### THAP1 binds to and regulates the *GusB* (MPS VII) gene encoding the GAG degrading enzyme β-glucuronidase

To mechanistically dissect the connection between THAP1 and GAG metabolism, we examined our RNAseq analyses (DIV2 and DIV2 & 4 common; Fig. 1F & Table S1). We identified a set of 42 genes (Table S3) relevant to GAG and proteoglycan metabolism that were differentially regulated in *Thap1^-/-^* cultures (Table S2). We focused our attention on genes that are (1) directly involved in GAG metabolism and (2) direct targets of THAP1. These criteria identified the *Gusb* gene encoding β-glucuronidase as a strong THAP1 candidate target. β-glucuronidase is an essential lysosomal enzyme that participates in the later steps of the catabolism of multiple species of GAG polysaccharides and proteoglycans. Recessively inherited loss-of-function mutations in *GUSB* causes the mucopolysaccharidosis Sly’s syndrome (MPS VII) (Heuer et al., 2001; Kubaski et al., 2017; Kyle et al., 1990; Paigen, 1989; Ray et al., 1999; Zielonka et al., 2017). Prior ChIPseq data from human K562 cells demonstrates that THAP1 is bound to the *GUSB* promoter (Fig. 4A; ENCODE dataset). We confirmed that THAP1 plays a critical role in the transcription of the *GusB* locus in the OL lineage (Fig. 4B-E). Quantitative chromatin immunoprecipitation (qChIP) analyses in DIV4 OL demonstrated that THAP1 is bound to the *GusB* promoter in the OL lineage (Fig. 4B). Consistent with that finding, *GusB* expression is significantly and profoundly reduced in *Thap1^-/-^*cultures (OPC through DIV4; Fig. 4E). We previously demonstrated that most THAP1 targets are also bound by YY1, a transcription factor known to regulate OL maturation, and that YY1 DNA binding is significantly reduced in *Thap1^-/-^* OLs (Yellajoshyula et al., 2017). Analyses of the *GusB* locus demonstrate that it is bound by YY1 in a THAP1-dependent manner (Fig. 4C). H3K4me3, an epigenetic modification associated with active transcription, is significantly reduced at the *GusB* locus in *Thap1^-/-^* OL, further connecting THAP1 loss to *GusB* function (Fig. 4D).

**Figure 4.**
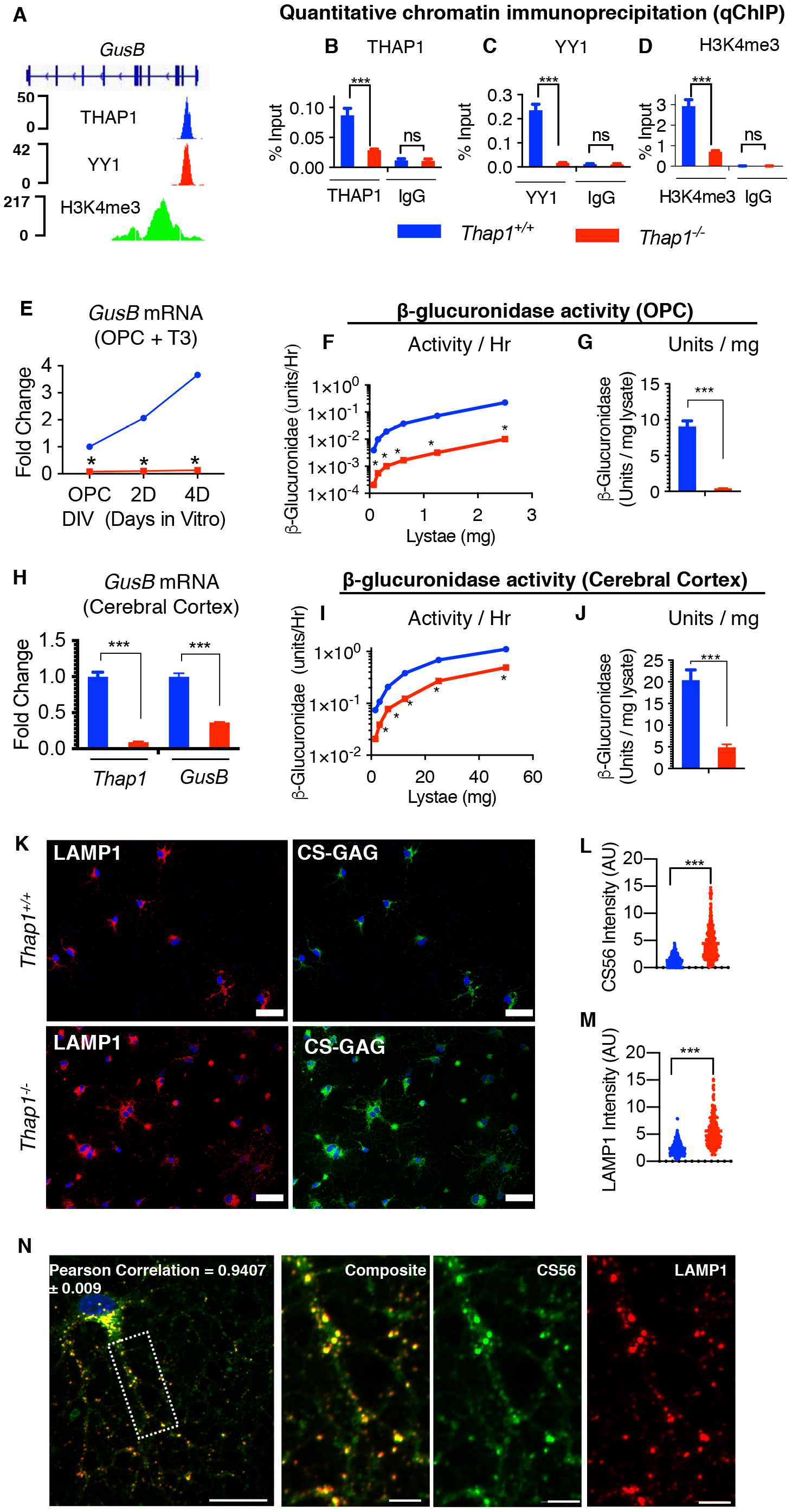
THAP1 directly binds to and regulates the *GusB* gene encoding the GAG-catabolizing lysosomal enzyme β-glucuronidase. (A) Genome browser track (using Integrative Genomics Viewer) showing CHIP-Seq signals for THAP1, YY1 and H34me3 at the *GusB* locus in K652 cells (ENCODE dataset). (B-D) Quantitative ChIP (qChIP) demonstrating THAP1 dependent binding of YY1 and H3K4me3 methylation at the *GusB* promoter region in OL (+T3 - DIV4). Binding, represented as % input (y-axis), is demonstrated for (B) THAP1 (x-axis) (*Thap1^+/+^* = 0.087% ± 0.008; *Thap1^-/-^* = 0.02% ± 0.001; t-test: t_(4)_ = 6.85; p = 0.02); (C) YY1 (x-axis) (*Thap1^+/+^* = 0.23% ± 0.017; *Thap1^-/-^* = 0.018% ± 0.0009; t-test: t_(2)_ = 12.85; p = 0.006); and (D) H3K4me3 (x-axis) (*Thap1^+/+^* = 2.93% ± 0.219; *Thap1^-/-^* = 0.707% ± 0.036; t-test: t_(2)_ = 10.04; p = 0.0098) and their respective isotype control IgG (Goat IgG or Rabbit IgG) at the *GusB* promoter region. Bar graph shows mean values ± SEM. Percentage input represents the final amount of immunoprecipitated chromatin/gene from corresponding *Thap1^+/+^* and *Thap1^-/-^* OL cells (Methods). (E-J) THAP1 loss significantly reduces *GusB* mRNA expression and β-glucuronidase enzyme activity in OLs and in CNS tissue. (E) *GusB* mRNA expression (qRT-PCR) from *Thap1^+/+^* and *Thap1^-/-^* differentiating OL (OPC, DIV2 & DIV4). *GusB* expression was normalized to *Rpl19* expression and is represented as fold change (y-axis) with respect to *Thap1^+/+^.* Line graphs show mean values ± SEM. (F-G) Deficient β-glucuronidase enzyme activity in *Thap1^-/-^* OPC. Line graph shows the mean value of β-glucuronidase activity (y-axis = Units/Hr) for (F) *Thap1^+/+^* and *Thap1^-/-^* OPC lysate (0 - 3 μg; x-axis) derived from fluorometry assay (Methods). Bar graph shows mean ± SEM values of normalized β-glucuronidase activity Units/Lysate (mg) for (G) OPC (*Thap1^+/+^* = 9.05 U / mg ± 0.803 ; *Thap1^-/-^* = 0.396 U / mg ± 0.038; t-test: t_(10)_ = 10.76; p < 0.0001). (H) *GusB* mRNA expression (qRT-PCR) from control (*Thap1^+/flx^)* and THAP1 N-CKO (*Thap1^flx/-^*; Nestin-*Cre^+^*) CNS (cerebral cortex, age P21). *GusB* expression was normalized to *Rpl19* expression and is represented as fold change (y-axis) with respect to control. Line graphs show mean values ± SEM. (I-J) Deficient β-glucuronidase enzyme activity in THAP1 N-CKO CNS. Line graph shows the mean value of β-glucuronidase activity (y-axis = Units / Hr) for (I) control (*Thap1^+/flx^)* and THAP1 N-CKO (*Thap1^flx/-^;* nestin-*Cre^+^*) cerebral cortex lysate (0 - 60 μg; x-axis) derived from fluorometry assay. (J) Bar graph shows mean ± SEM values of normalized β-glucuronidase activity Units/Lysate (mg) for (J) CNS (Control = 20.43 U / mg ± 2.35 ; N-CKO = 4.90 U / mg ± 0.68; t-test: t_(10)_ = 6.34; p < 0.0001). 2-way ANOVA demonstrates a main effect of genotype (P<0.01) for (E,H) *GusB* mRNA expression and (F,J) β-glucuronidase activity in (E,F) OPC or (H,I) cerebral cortex. (*) represents time points where significant differences exist using post hoc Sidak’s multiple comparison test. (K) Representative images (scale bar 50 μm) and (L-M) corresponding quantification of *Thap1^+/+^* and *Thap1^-/-^* OL cells (DIV4) immunostained under conditions of membrane permeabilization for CS-56 (CS-GAG; *Thap1^+/+^* = 1.34 ± .072; *Thap1^-/-^* = 4.44 ± 0.16; t-test: t_(530)_ = 14.51; p < 0.0001) or LAMP1 (*Thap1^+/+^* = 2.36 AU ± 0.082; *Thap1^-/-^* = 5.57 AU ± 0.20; t-test: t_(412)_ = 15.27; p < 0.0001). Each data point in the scatter plot represents CS-56 or LAMP1 staining intensity for individual cells. (N) Representative images (scale bar 20 μm) of CS-GAG (CS-56) co-localization with lysosomes (LAMP1) in OL (DIV4). Pearson’s coefficient value (0.94 ± 0.009) shows ~ 95% colocalization (R = 0.94 ± 0.009) (Fig. S4D).

Consistent with significantly reduced *GusB* expression in *Thap1^-/-^* OL (Fig. 4E), we also find reduced β-glucuronidase enzymatic activity in cell culture and CNS tissue, as measured by fluorometry (MUG assay). In *Thap1^+/+^* cultures, β-glucuronidase activity was comparable in OPC and OL (DIV4) (Fig. S4A-B). *Thap1^-/-^* OLs exhibited a nearly 20-fold decrease of β-glucuronidase activity (Fig. 4F-G). In *Thap1^-/-^* cortical tissue, *GusB* mRNA was decreased by ~80% (Fig. 4H) and β-glucuronidase activity was reduced by ~75% (Fig. 4I-J). Several other brain regions assessed (cortex, striatum, corpus callosum and cerebellum) showed similar reductions in β-glucuronidase activity (Fig. S4C). β-Glucuronidase is a lysosomal enzyme, suggesting that loss of its activity should cause lysosomal accumulation of CS-GAGs. Indeed, assessing intracellular CS-GAG by staining permeabilized cells demonstrated significantly higher levels in *Thap1^-/-^* OL cultures (Fig. 4K-M). Co-labeling cells for CS-GAG and the pan-lysosomal marker LAMP1 confirmed that the intracellular CS-GAG accumulation is almost exclusively localized to lysosomes (Fig. 4N, S4D). Considered together with the effects of THAP1 at the *GusB* locus, these data indicate a direct and essential role for THAP1 in regulating levels of CNS β-glucuronidase.

### *GUSB* overexpression rescues THAP1 mutant GAG homeostatic and oligodendroglial maturation defect *in vitro* and *in vivo*

To formally test the *necessity* of reduced β-glucuronidase for THAP1-related GAG homeostatic and oligodendroglial lineage defects, we pursued rescue studies with a *GUSB* transgene (Tg^GUS;^ (Kyle et al., 1990)). Prior work demonstrated that the Tg^GUS^ does not produce a phenotype when in the hemizygous state on a wild type background (Kyle et al., 1990). We ingressed Tg^GUS^ onto the CNS conditional *Thap1^-/-^* background. We used these mice to derive OPCs from SVZ derived neural stem cells (as previously described (Yellajoshyula et al., 2017)) from *Thap1^+/+^* and *Thap1^-/-^* with or without the Tg^GUS^ transgene. We confirmed that the Tg^GUS^ transgene significantly increased β-glucuronidase activity in *Thap1^-/-^* OPCs (Fig. 5A) and that *Thap1^-/-^;Tg^GUS^* OPCs normally express the OPC-specific markers OLIG2 and CSPG4 (NG2) (Fig. 5B).

**Fig. 5.**
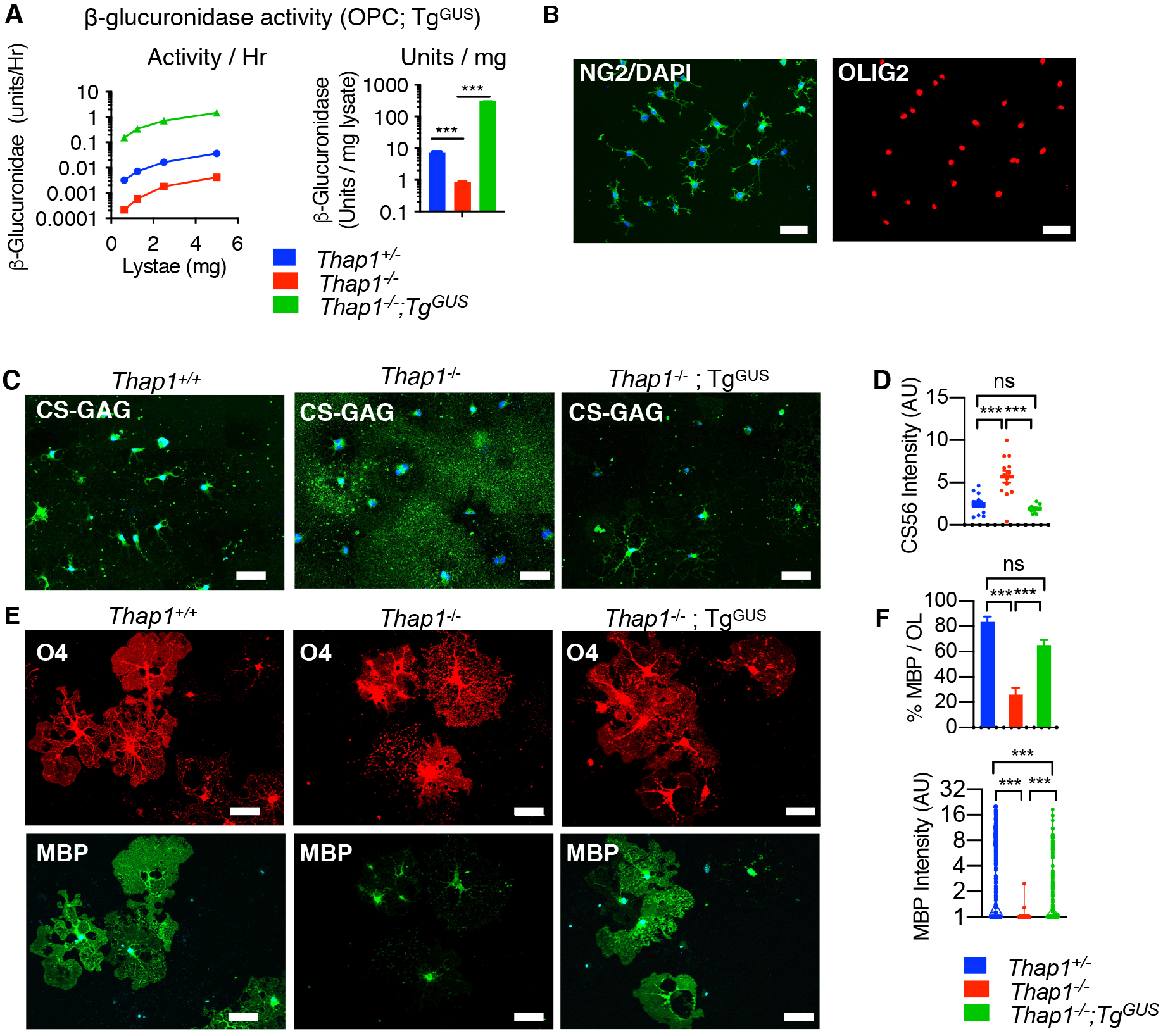
β-glucuronidase overexpression rescues the *Thap1* null GAG homeostatic and OL maturation defects. (A) β-glucuronidase activity in OPC lysate from *Thap1^+/+^, Thap1^-/-^* and *Thap1^-/-^*;Tg^GUS^ genotypes (mean ± SEM of normalized β-glucuronidase activity; *Thap1^+/+^* = 7.48 U / mg ± 0.193; *Thap1^-/-^* = 0.87 U / mg ± 1.29; *Thap1^-/-^;* Tg^GUS^ = 304.1 U / mg ± 1.6; One-way ANOVA F_(2,149972)_ = 34628.96, p<0.0001, Dunnett’s multiple comparisons test: adjusted p value < 0.0001). (B) Representative images (scale bar 50 μm) of *Thap1^-/-^*;Tg^GUS^ OPC expressing NG2 (CSPG4) and pan-OL lineage marker OLIG2. (C-D) β-glucuronidase overexpression prevents CS-GAG accumulation in *Thap1^-/-^* OL. (C) Representative images (scale bar 50 μm) and (D) corresponding quantification for *Thap1^+/+^,Thap1^-/-^* and *Thap1^-/-^*;Tg^GUS^ OL (+T3 - DIV5) immunostained with CS-56 (CS-GAG) under non-permeabilized conditions. Each data point in the scatter plot represents CS-56 staining intensity for individual cells normalized to area. *Thap1^+/+^* = 2.46 AU ± 0.35; *Thap1^-/-^* = 5.7 AU ± 0.67; *Thap1^-/-^*;Tg^GUS^ = 1.909 AU ± 0.13; One-way ANOVA F_(2,34)_ = 19.95, p<0.0001, Dunnett’s multiple comparisons test: adjusted p value < 0.0001. (E-F) β-glucuronidase overexpression rescues maturation defects in *Thap1^-/-^* OL. (E) Representative images (scale bar 50 μm) for *Thap1^+/+^* and *Thap1^-/-^* and *Thap1^-/-^*;Tg^GUS^ OL differentiated for 5 days (+T3). Cultures stained with O4, MBP, & DAPI. (F) Quantification of the % of O4+ cells expressing MBP and intensity of MBP staining. (F, upper graph) Percentage of O4+ cells expressing MBP. Bar graph shows mean ± SEM values. y-axis; *Thap1^+/+^* = 83.48% ± 5.08 ; *Thap1^-/-^* = 26.17% ± 5.33; *Thap1^-/-^*;Tg^GUS^ = 65.18 ± 4.09; One-way ANOVA F_(2,49)_ = 23.27, p<0.0001, Dunnett’s multiple comparisons test: adjusted p value < 0.0001. (F, lower graph) Mean intensity of MBP staining normalized to background. Each data point in the scatter plot represents MBP+ staining intensity for individual cells. *Thap1^+/+^* = 4.81 AU ± 0.24; *Thap1^-/-^* = 1.005 ± 0.004; *Thap1^-/-^*;Tg^GUS^ = 1.933 ± 0.128; One-way ANOVA F_(2,1195)_ = 127.8, p<0.0001, Dunnett’s multiple comparisons test: adjusted p value < 0.0001.

We began assessing the effects of increasing β-glucuronidase by measuring CS-GAG levels in *Thap1^-/-^* OPCs. Immunostaining *Thap1^-/-^;Tg^GUS^* OL (+T3-DIV4) for CS-GAG under non-permeabilizing conditions demonstrated that the Tg^GUS^ transgene completely reversed the CS-GAG accumulation phenotype (Fig. 5C-D). At DIV4, *Thap1^-/-^*;Tg^GUS^ cultures also exhibited a significantly greater number of myelinating OL (%MBP/OL;Fig. 5E-F) and increased MBP staining (MBP+ intensity; Fig. 5E-F) compared to *Thap1^-/-^* OL. Having confirmed *in vitro* rescue we next tested the ability of Tg^GUS^ transgene to rescue THAP1-related myelination defects *in vivo.* We confirmed that the Tg^GUS^ allele increases β-glucuronidase activity in all CNS regions tested (Fig. S5). Using electron microscopy we assessed 1) density of myelinated axons; 2) myelin thickness; and 3) size of myelinated axons. We performed these analyses in P21 tissue from the genu of corpus callosum tissue from CNS conditional knockout *Thap1* (nestin-cre; “N-CKO”) or control (*Thap1^+/+^)* mice with or without the Tg^GUS^ transgene (Fig. 6 A-F). N-CKO;Tg^GUS^ mice exhibited a 2-fold increase in the density of myelinated axons compared to N-CKO mice without the transgene (Fig. 6 A-B). Tg^GUS^ did not fully restore myelinated axon density to control levels, however. The abnormally high g-ratio characteristic of N-CKO mice, in contrast, was rescued completely by restoration of β-glucuronidase (Fig. 6 A,C-D,F). N-CKO mice exhibit a modest but significant increase in the size of the myelinated axons, which was also rescued by the Tg^GUS^ transgene (Fig. 6 A,E). These findings establish β-glucuronidase activity as a key mediator of THAP1-related CNS pathology.

**Fig. 6.**
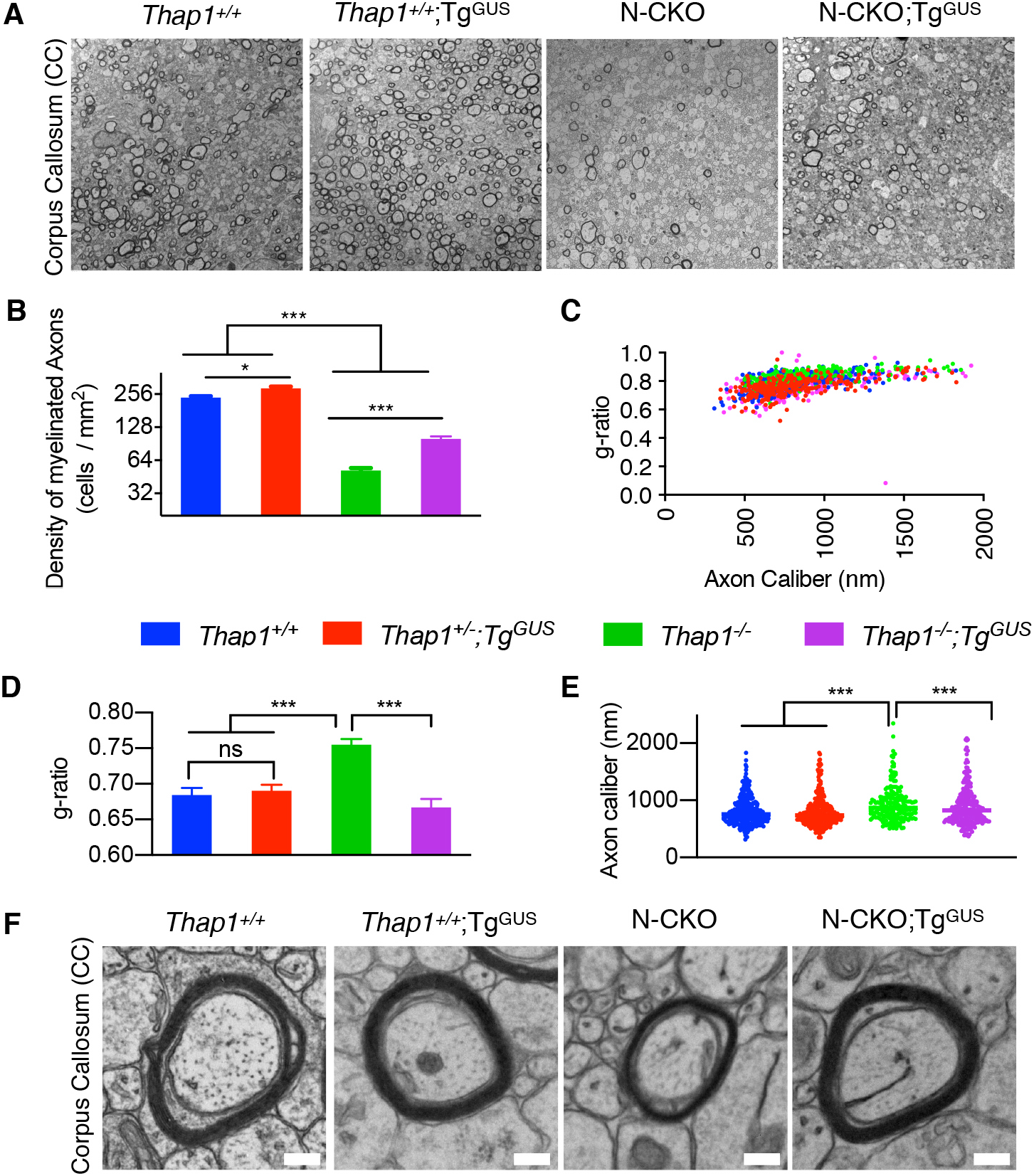
β-glucuronidase overexpression rescues the myelination defect in the THAP1 null CNS. (A) Representative transmission EM images (scale bar 4 μm) of the genu of corpus callosum at P21 from control (*Thap1^+/+^;* nestin-*Cre^-^*), control-Tg^GUS^ (*Thap1^+/+^;* nestin- *Cre^-^),* N-CKO (*Thap1^flx/-^;* nestin-*Cre^+^*) and N-CKO-Tg^GUS^ (*Thap1*f^x/-^; nestin-*Cre^+^*; Tg^GUS^). (B-E) Quantification of EM images (N = 3 per genotype) demonstrates that β-glucuronidase overexpression (N-CKO; Tg^GUS^) significantly increases myelination in the THAP1 null CNS. (B) Density of myelinated axons (number of axons / mm^2^). Mean ± SEM; Control = 238.09 ± 12.58, Control;Tg^GUS^ = 289.85 ± 10.36; N-CKO = 51.74 ± 2.70; N-CKO;Tg^GUS^ 100.71 ± 4.790; One-way ANOVA F_(3,6)_ = 182.7, p<0.0001, Dunnett’s multiple comparisons test: adjusted p value < 0.0001. (C-D) g-ratio represented in relation to axon caliber (C) and by genotype (D, mean ± SEM; Control = 0.684 ± 0.009, Control;Tg^GUS^ = 0.690 ± 0.0087; N-CKO = 0.755 ± 0.0082; N-CKO;Tg^GUS^ = 0.666 ± 0.0119; One-way ANOVA F_(3,8)_ = 15.68, p = 0.001, Dunnett’s multiple comparisons test: adjusted p value < 0.004). (E) Axon caliber (each data point represents axon caliber (nm) of individual axons; Control = 813.2 nm ± 14.76, Control;Tg^GUS^ = 825.9 nm ± 19.54; N-CKO = 935.0 ± 21.42; N-CKO;Tg^GUS^ = 897.4 ± 21.08; One-way ANOVA F_(3,1047)_ = 8. 613, p = 0.001, Dunnett’s multiple comparisons test: adjusted p value < 0.004). (F) Representative EM images (scale bar 200 nm) demonstrating rescue of myelin thickness (g-ratio) observed in N-CKO with β-glucuronidase overexpression (N-CKO; Tg^GUS^).

## DISCUSSION

Our studies establish THAP1 as a transcriptional regulator of OL maturation via its control of GAG-metabolism. We demonstrate that THAP1 directly binds to and regulates the *GusB* gene encoding β-glucuronidase, a lysosomal GAG-degrading enzyme. THAP1 loss dramatically decreases *GusB* mRNA and β-glucuronidase enzymatic activity, causing CS-GAG lysosomal accumulation and increasing secretion. Multiple lines of evidence support CS-GAG accumulation as the key mechanism responsible for delaying maturation of the *Thap1^-/-^* OL lineage. Preventing abnormal accumulation of GAGs rescues the *Thap1^-/-^* OL maturation defect whereas administering GAG species that accumulate in *Thap1^-/-^* cultures (C-4S, C-6S) impairs myelination of control OLs. Moreover, overexpressing *GUSB* rescues GAG accumulation and OL maturation, and ameliorates CNS myelination delay in *Thap1^-/-^* mice. Considered together, our findings identify a novel cell autonomous mechanism of OL lineage regulation and provide insight into the molecular pathophysiology of THAP1 loss-of-function, the cause of DYT6 dystonia.

The role of CS-GAGs (and CSPGs) in regulating OL differentiation has been well studied (Harlow and Macklin, 2014; Karus et al., 2016; Keough et al., 2016; Kuboyama et al., 2017; Lau et al., 2012; Pu et al., 2018; Siebert and Osterhout, 2011; Sloane et al., 2010). CSPGs have been demonstrated to inhibit OL differentiation through their interactions with the OPC resident receptor-type protein-tyrosine phosphatase (RPTP) subtypes, RPTPζ and RPTPσ, to suppress the activation of Rho-associated kinase (ROCK), which is antagonistic to OL differentiation (Kuboyama et al., 2016; Pendleton et al., 2013). Studies exploring the cellular source of these signals have largely focused on astrocytes (Asher et al., 2000; Feliu et al., 2020; Laabs et al., 2007; McKeon et al., 1999; Tang et al., 2003). Here we demonstrate that the OL lineage itself is a functionally important source of CS-GAGs that is auto-inhibitory to its development. Several prior and recent studies support our hypothesis that include the findings of CSPG production from OL cultures (Courel et al., 1998; Garwood et al., 2004; Karus et al., 2016; Niederost et al., 1999; Probstmeier et al., 2000; Szuchet et al., 2000; Yim et al., 1993) and the application of ChABC, inhibiting CSPG production or inactivation of CSPG receptor PTPRZ in purified OL cultures promote their differentiation (Karus et al., 2016; Kuboyama et al., 2015). Our mass spectrometry analyses indicate that within the OL lineage, CS-GAGs are uniquely secreted by OPCs. OL lineage progression is accompanied by a precipitous drop in CS-GAG levels. This drop appears to be an essential step in OL maturation. Our studies suggest that the drop in CS-GAGs is not likely from increased catabolism, as the β-glucuronidase activity remans comparable through maturation (Fig. S4). Rather, the expression of genes involved in CS-GAG production (identified from GO terms) decrease through OL differentiation.

We utilized LC/MS studies to analyze three GAGs (CS, HS and HA) with well-established roles in CNS development. We discovered that C-4S is the major GAG species secreted (~ 85% of total CS-GAG) by the OL lineage (Fig. 3). Differentially sulfated CS-GAGs mediate unique biological activities (Properzi et al., 2005). The discovery of C-4S production within OPCs and its auto-inhibitory activity on these cells raises important questions about CS-GAG metabolism in normal development and in the context of brain disease and repair. Despite the biological activity conferred by the GAGs in their free form and as proteoglycans, there have limited reports exploring GAG metabolism or secretion in the OL lineage. Rather, secreted proteoglycans (e.g., aggrecan family proteins) have been the major focus of studies exploring GAG/ proteoglycan significance in injury and development.

How does THAP1 LOF cause GAG accumulation? We demonstrate that *GusB* is the critical downstream effector for this phenotype. GusB encodes β-glucuronidase, a lysosomal enzyme that catalyzes the terminal stages of GAG catabolism by hydrolyzing glucuronide or glucuronic acid from multiple GAG species (CS-, HS- and HA-GAGs) (Heuer et al., 2001; Ray et al., 1999). Complete loss of *GUSB* causes Sly syndrome (MPS VII) (Beaudet et al., 1974; Sly et al., 1974), a lysosomal storage disease with marked GAG elevations (Heuer et al., 2001; Kubaski et al., 2017; Ray et al., 1999), lysosomal abnormalities (Bayo-Puxan et al., 2018; Paigen, 1989; Vogler et al., 1990) and multi-organ defects including neurodevelopmental, cognitive and motor dysfunction (Montano et al., 2016; Sands, 2014; Tomatsu et al., 2009). Importantly and consistent with our findings, MPS VII patients and mouse models (*GusB*^mps^) (Kumar et al., 2014) exhibit reductions in CNS myelination and altered expression of myelin component genes (Parente et al., 2012). Myelin abnormalities also occur in other MPS syndromes (Reichert et al., 2016; Seto et al., 2001). Most studies of MPS VII pathogenesis in the CNS have focused on *GusB* function and regulation in *neurons* (Bayo-Puxan et al., 2018; Levy et al., 1996; Parente et al., 2012; Parente et al., 2016; Sly and Vogler, 1997, 2002; Vogler et al., 1998). In contrast, our studies constitute the first effort defining the role of *GusB* in the glial lineage. *Klotho*, which codes for a type I membrane protein with glucuronidase activity and is related to β-glucuronidase has been shown to promote the maturation of OL lineage (Chen et al., 2013; Zeldich et al., 2015). However, *Klotho* is not a resident of lysosomes, has not been implicated in any MPS, and it is unknown if it plays any role GAG metabolism. Restoration of β-glucuronidase restores GAG homeostasis in the OL lineage, rescues *in vitro* maturation of *Thap1^-/-^* OLs, and significantly improves *Thap1^-/-^* CNS myelination. These observations establish lysosomal catabolism of GAGs as an essential element of THAP1 CNS function.

Most DYT6 dystonia mutations in *THAP1* are predicted to cause LOF, including early stop mutations (Blanchard et al., 2011; Bragg et al., 2011). Our prior studies demonstrate that THAP1 has a prominent role in CNS myelination during juvenile developmental, the period when dystonia emerges in most DYT6 subjects (Yellajoshyula et al., 2017). These observations highlight a critical role for lysosomal GAG catabolism during CNS maturation, and implicate this pathway in the neurodevelopmental disease DYT6 dystonia. While the experiments presented here focus exclusively on GAG effects within and on the OL lineage, considerable data suggest that GAG abnormalities impact neural plasticity and function (Lin et al., 2008; Smith et al., 2015). These non-cell autonomous effects, the impact of potentially abnormal GAG metabolism within neurons themselves, and the impact of delayed OL maturation on neural function are important future directions in the study of DYT6 pathogenesis, the role of ECM abnormalities in brain injury and disease, and fundamental understanding of neuronal-glial interactions.

## MATERIALS & METHODS

### Generation and Maintenance of Mice

Animal research was conducted in accord with the NIH laboratory animal care guidelines and with the Institutional Animal Care and Use Committee (IACUC) at the University of Michigan. *Thap1* floxed mice generation, characterization and genotyping have been previously described (Yellajoshyula et al., 2017). Nestin-*Cre*+ and Tg^GUS^ (Stock # 006558) was purchased from Jackson Laboratory (Bar Harbor, ME, USA). Olig2-*Cre* mice were kindly provided by Dr. Roman Giger. The breeding strategy used to generate derive all primary OPC cells and conditional null mice was as follows: *Thap1^+/–^, Cre^+^,Tg^GUS^ × Thap1^flox/flox^.* This breeding strategy produced the following offspring ± Tg^GUS^: *Thap1^flox/+^; Thap1^flox/–^; Thap1^flox/+^, Cre^+^;* and *Thap1^flx/–^, Cre^+^* (CKO). Age and sex-matched littermate mice were used for all experiments. Primers used for genotyping in this study (*Thap1, Cre* & *Tg^GUS^)* are listed in Table S4.

### RNA extraction, qRT-PCR and RNAseq

Total RNA extraction from OL cultures for qRT-PCR and RNAseq analysis was done using NucleoSpin® RNA (Takara) and Trizol (Thermofisher) for mouse cerebral cortex as per manufacturers instructions. cDNA synthesis from total RNA was done using MMLV Reverse Transcriptase (Takara) as per manufacturers instructions. Quantitative real time PCR (qRT–PCR) was performed with the StepOnePlus System (ABI) and 2x SYBR Power Mix (ABI). RNAseq was performed on Illumina HISeq2500 (Illumina) platform at the neurogenetics laboratory, NIA. RNAseq libraries were made using the TruSeq Stranded Total RNA Library prep kit with rRNA removal mix Gold (Illumina, 20020598). Libraries were quantified and normalized using the KAPA Library Quantification kit for Illumina Platforms (KAPA Biosystems, KR0405) and sequenced on an Illumina HiSeq 2500 using HiSeq SBS kit v4 250 cycle kit (Illumina, FC-401-4003). A standard Illumina pipeline was used to generate fastq files. RNAseq Analyses was performed using DESEQ.

### Immunostaining

Immunofluorescence was done as previously described (Yellajoshyula et al., 2017) for the following antibodies: Rat anti-MBP (Millipore, Cat# MAB386), Mouse anti-O4 (kindly provided by Dr. Roman Giger), Goat anti-OLIG2 (R&D, AF2418), Rabbit anti-NG2 (Millipore, AB5320) and Rat anti-LAMP1 (Santacruz, sc-19992). Staining for CS and HA GAGs with Mouse anti-CS-56 (Thermofisher, MA1-83055) and Biotinylated HABP (Millipore, 385911) was performed in PBS + 1% BSA with no detergent, unless mentioned otherwise. Quantification of staining intensity (normalized for area) was performed using imageJ analysis by tracing individual cells for CS-56, and using intensity of the entire image for HABP (to account for amorphous staining of HA-GAGs under non-permeabilizing conditions).

### Electron Microscopy

EM was performed as previously described (Yellajoshyula et al., 2017). Briefly, P21 mice were anesthetized and perfused with EM perfusion solution (3% paraformaldehyde, 2.5% glutaraldehyde in 0.1M phosphate buffer). Following perfusion, brains were dissected and postfixed at 4°C overnight in perfusion solution. Genu of CC was dissected, processed and sectioned at EM core facility, Emory University. All EM images were acquired using JEOL JSM 1400 at the University of Michigan, MIL core services. Axon caliber and g-ratio (ratio of the inner axonal diameter to total (including myelin) outer diameter) were calculated as previously described (Winters et al., 2011; Yellajoshyula et al., 2017).

### Derivation of OPC from NSC cells

OPC used in this study were derived from primary NSC isolated from the sub ventricular zone (SVZ) of P7 mouse pups as previously described (Yellajoshyula et al., 2017). Routine propagation of the OPC was done as a monolayer on poly-L-ornithine (0.1mg/ ml) and laminin (5μg/ml) coated dishes in OPC expansion media (SATO medium supplemented with 20μg/ml PDGF and FGF2; (Yellajoshyula et al., 2017)) or differentiated on poly-L-ornithine (0.1mg/ml) and poly-D-lysine (0.1mg/ml) coated coverslips or dishes in oligodendrocyte differentiation medium (SATO media with 40μg/ ml T3) (Yellajoshyula et al., 2017).

### β-glucuronidase activity assay - MUG assay

Tissue or cell pellets were lysed in MUG lysis buffer (0.2M Sodium acetate buffer, pH 4.5;10 mM EDTA; 0.1% Trixton-X-100) to obtain total cell homogenate. Equal concentrations of serially diluted lysate (in the range of 0-60 μg/ml) from control and *Thap1* cKO genotypes was incubated with 10 mM 4-methylumbelliferyl β-D-glucuronide (MUG, Sigma) in 0.2M Sodium acetate buffer, pH 4.5 at 37°C for one hour. The reaction was stopped with equivalent volume of 0.2M sodium citrate followed by which the fluorescence (excitation = 360nm; emission = 460nm) was measured using Biotek Synergy HT microplate reader. Based on fluorescence reading from purified β-glucuronidase standard (Sigma), we determined the β-glucuronidase activity / hour and β-glucuronidase activity / mg of the lysate.

### Quantitative Chromatin immunoprecipitation (qCHIP)

qCHIP was performed as previously described (Yellajoshyula et al., 2011). Sheared chromatin (sonicated to 200–500 bp) from 2 × 10^6^ mouse OL cells was incubated with 2.5μg of Goat THAP1 (sc-98174, Santacruz), Rabbit YY1 (sc-98174, Santacruz), Rabbit H3K4me3 (sc-98174, Activemotif), normalized Goat IgG (Santacruz), or normalized Rabbit IgG (Santacruz) using Dynabeads (Invitrogen). After washing, elution and crosslink reversal, DNA from each ChIP sample and the corresponding input sample was purified and analyzed further using qPCR. Each ChIP sample and a range of dilutions of the corresponding input sample (0.01 – 2% input) were quantitatively analyzed with gene-specific primers using the StepOnePlus System (ABI) and SYBR qPCR Powermix (ABI).

### Gene Ontology enrichment analysis

Candidate THAP1-dependent DEGs identified from RNAseq catalysis were analyzed for enrichment of biological pathways by gene ontology (GO) enrichment analyses using the following web based applications: GENEONTLOGY http://geneontology.org/docs/go-enrichment-analysis/ and DAVID https://david.ncifcrf.gov/home.jsp. Top GO enriched (p <0.01 and > 2 fold enrichment) for GO-Biological processes and GO-Molecular function was used for this study.

## Statistics

All data are reported as mean ± SEM. All statistical tests reported (Student’s t-tests, One-way or two-way ANOVAs) were performed using Graphpad Prism software (V9).

## Data & Software Availability

The GEO accession number for CHIP-seq data used in manuscript for THAP1 is GSM803408, YY1 is GSM803446 and H3K4me3 is GSM788087.

## ACKNOWLEDGEMENTS

We thank Haoran Huang and Stephanie Mrowczynski for technical assistance. We thank Cathy Collins and Jay Li for critical reading of manuscript. We are grateful to the staff of UCSD Glycoanalytics Core for GRIL LC/MS analysis. We thank Dr. Herbert Geller for helpful suggestions. We thank the staff of University of Michigan’s Core Facilities (DNA Sequencing Core, Microscopy and Image Analysis Laboratory and Unit of Laboratory Animal Medicine) and Hong Yi (EM Core, Emory University). This research was supported in part by the following grants: to WTD (7R56NS109227-02 NINDS).

