## Supplemental Data for "THAP1 Modulates Oligodendrocyte Maturation by Regulating ECM Degradation in Lysosomes"

#### Fig. S1. Related to Figure 1

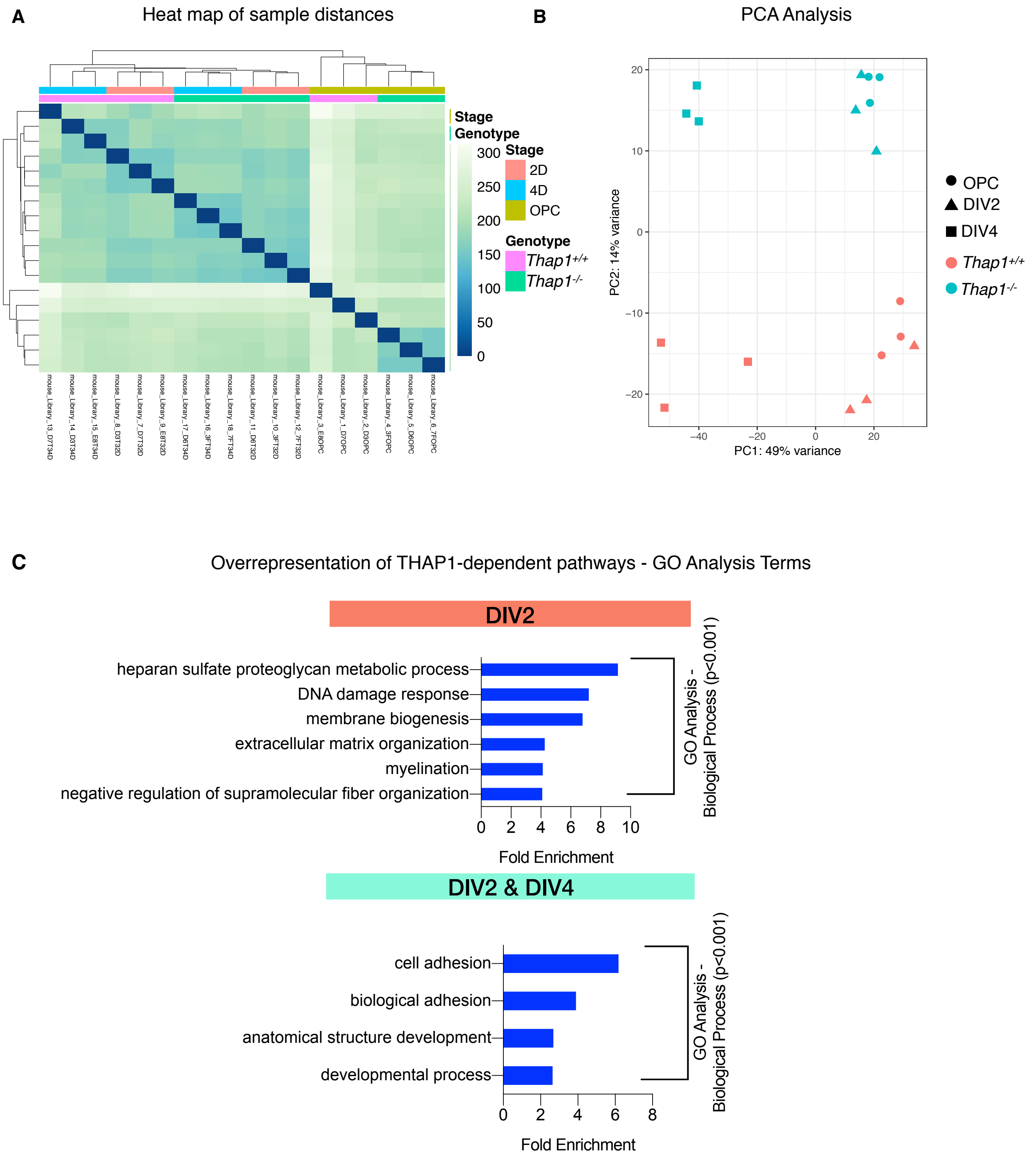

### Figure S2 - related to Figure 2

A

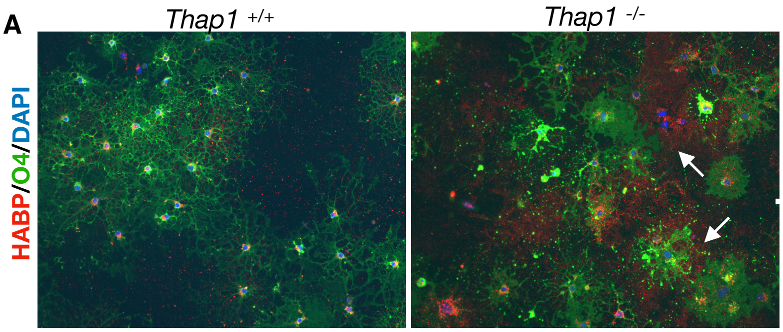


B

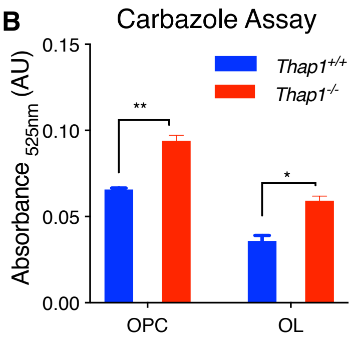


**C** Secreted GAG / Media (μg/mg)

| | OPC | | OL | |
|---|---|---|---|---|
| | *Thap1*+/+ | *Thap1*-/- | *Thap1*+/+ | *Thap1*-/- |
| CS | 73.53 | 134.07 | 9.40 | 23.56 |
| HS | 3.67 | 7.55 | 0.44 | 1.01 |
| HA | 1.40 | 2.61 | 0.86 | 2.42 |

**D** GAG / Homogenate (μg/mg)

| | OPC | | OL | |
|---|---|---|---|---|
| | *Thap1*+/+ | *Thap1*-/- | *Thap1*+/+ | *Thap1*-/- |
| CS | 151.64 | 405.42 | 169.35 | 550.80 |
| HS | 46.85 | 171.20 | 79.74 | 121.03 |
| HA | 161.77 | 186.62 | 290.16 | 448.82 |

**E** Secreted Chondroitin Sulfate GAG / Media (μg/mg)

| | OPC | | OL | |
|---|---|---|---|---|
| | *Thap1*+/+ | *Thap1*-/- | *Thap1*+/+ | *Thap1*-/- |
| C-4S | 63.15 | 123.88 | 7.77 | 19.76 |
| C-6S | 7.44 | 6.34 | 1.00 | 2.48 |
| C-2,6S | 0.82 | 1.17 | 0.13 | 0.44 |
| C-4,6S | 0.79 | 1.05 | 0.09 | 0.15 |

**F** Chondroitin Sulfate GAG / Homogenate (μg/mg)

| | OPC | | OL | |
|---|---|---|---|---|
| | *Thap1*+/+ | *Thap1*-/- | *Thap1*+/+ | *Thap1*-/- |
| C-4S | 134.74 | 376.49 | 150.48 | 472.26 |
| C-6S | 13.62 | 13.40 | 15.21 | 48.29 |
| C-2,6S | 0.25 | 1.72 | 0.28 | 10.14 |
| C-4,6S | 2.09 | 5.00 | 2.33 | 5.22 |

#### Figure S3 - related to Figure 3

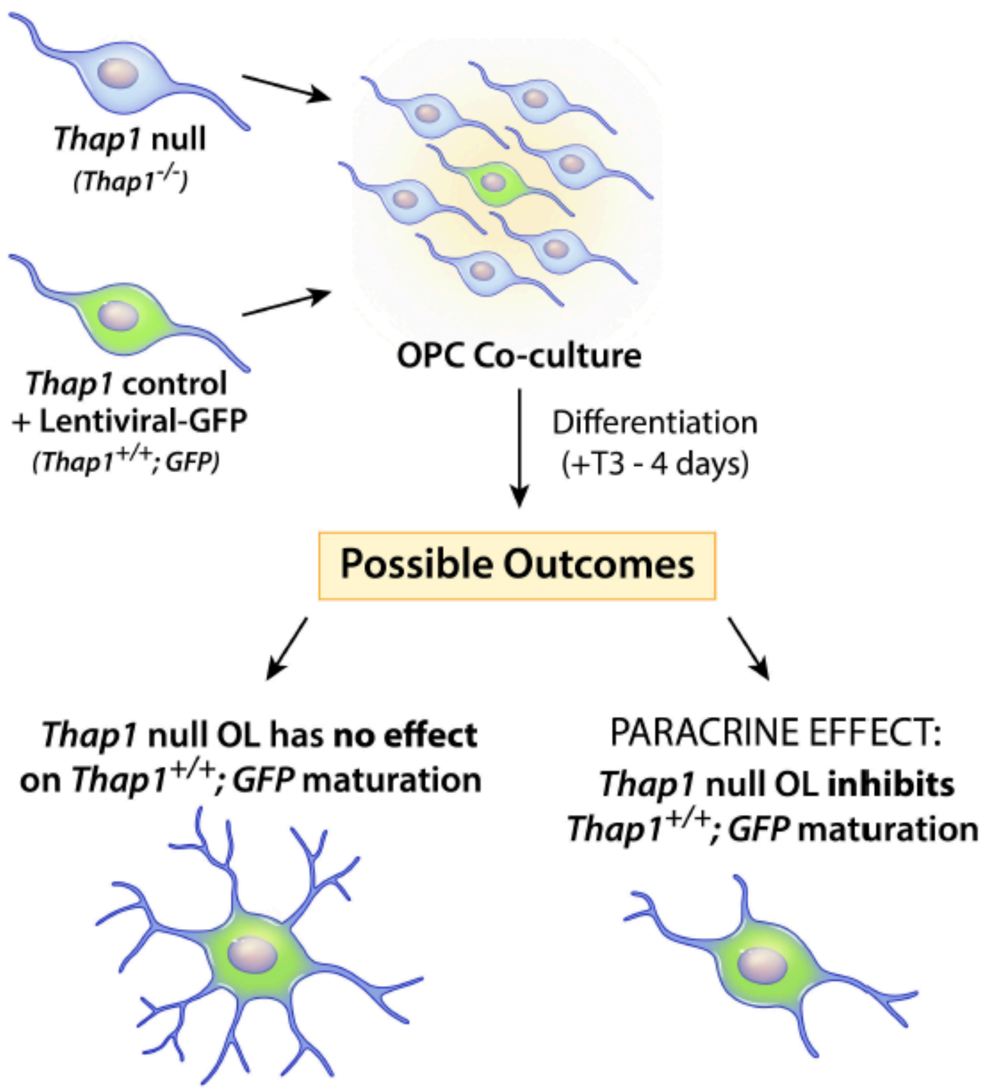

**Figure S4 - related to Figure 4**

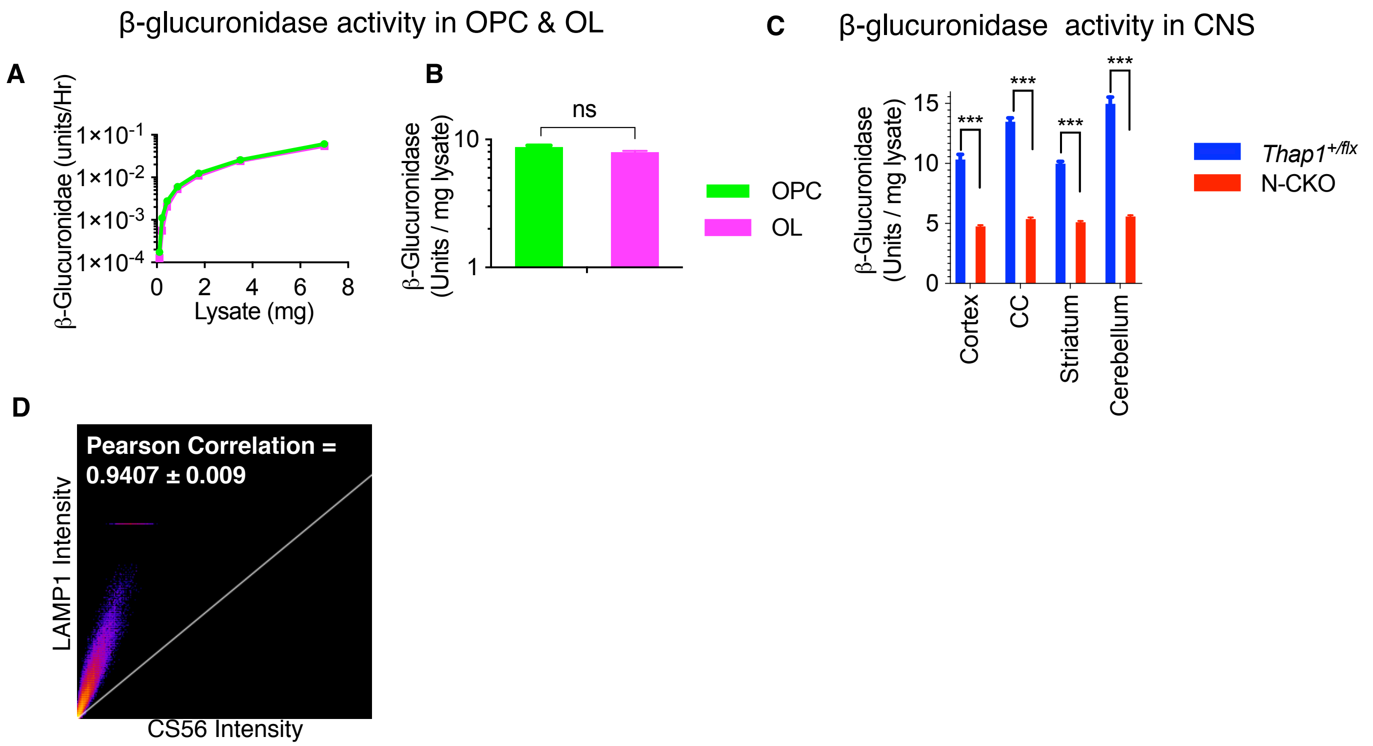

**Figure S5- related to Figure 5**

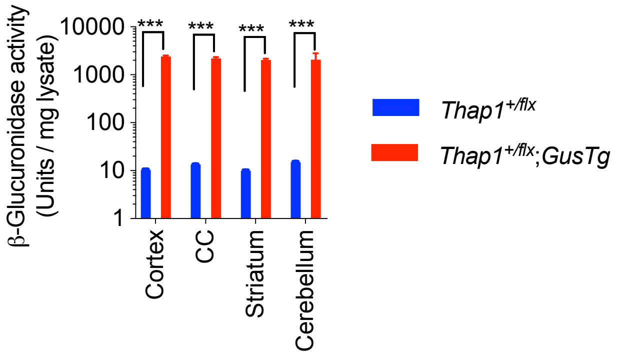

#### SUPPLEMENTAL FIGURE LEGENDS

**Figure S1 related figure 1. THAP1 loss disrupts the extracellular matrix transcriptome**

(A) Heatmap for sample distances and (B) principal component analysis (PCA) showing the variance in the samples used for RNAseq analysis: 3 clonal lines for each genotype (*Thap1*$^{+/+}$ and *Thap1*$^{-/-}$) from 3 stages of OL cultures: OPC, DIV2 & DIV4. (C) *Thap1*$^{+/+}$ vs *Thap1*$^{-/-}$ differentially expressed genes (DEG) selective for either DIV2 (703) or present at both DIV2 & DIV4 (277) used for enrichment analysis of GO (Gene Ontology) terms to identify overrepresented biological pathways. Graphs show the most significantly overrepresented GO terms (p<0.001, sorted as fold overrepresented) in OL from DIV2 (GENEONTOLOGY, GO - Biological Process), and from the DIV2 & DIV4 overlapping group (DAVID, GO - GO - Biological process)

**Figure S2 related figure 2. *Thap1* null oligodendrocytes accumulate and secrete excess glycosaminoglycans**

(A) Representative images demonstrating accumulation of HA-GAG (stained using hyaluronan binding protein, HABP; red) in *Thap1*$^{-/-}$ OL cultures (+T3-DIV4). Arrows indicate surrounding regions of select OL with HA-GAG accumulation. (B) Carbazole assay (demonstrating GAG content) of total cell homogenate extracted from *Thap1*$^{+/+}$ and *Thap1*$^{-/-}$ OPC and OL (+T3-DIV4). Absorbance at 525 nm for OPC; *Thap1*$^{+/+}$ = 0.066 AU ± 0.0006; *Thap1*$^{-/-}$ = 0.094 AU ± 0.0022; t-test: $t_{(2)}$ = 12.70; p = 0.006 & OL - *Thap1*$^{+/+}$ = 0.036 AU ± 0.0023; *Thap1*$^{-/-}$ = 0.059 AU ± 0.0019; t-test: $t_{(2)}$ = 7.96; p = 0.015. Table displaying the estimated amount of (C-D) CS, HS and HA GAGs (μg/mg) and (E-F) composition of mono (C-4S, C-6S) and bi-sulfated (C-2,6S and C-4,6S) CS-GAGs (μg/mg) (C,E) secreted in the media or (D,F) from cell homogenate from *Thap1*$^{+/+}$ and *Thap1*$^{-/-}$ OPC and OL (+T3-DIV4) using GRIL LC/MS analysis.

**Figure S3 related figure 3. Excess glycosaminoglycan secretion by *Thap1* null cells impairs oligodendroglial maturation via a non-cell autonomous mechanism**

Schematic illustrating co-culture experimental paradigm to test for paracrine effects of GAGs secreted by the OL lineage. *Thap1* null (*Thap1*$^{-/-}$) OL were co-cultured with LV-GFP labelled control (*Thap1*$^{+/+}$*; GFP*) OPC at 10:1 ratio in differentiation media for four days (+T3). The two possible outcomes from this experimental paradigm are depicted in the illustration.

**Figure S4 related figure 4. β-glucuronidase activity during oligodendrocyte differentiation.**

(A) β-glucuronidase activity (Units / Hr) for *Thap1*$^{+/+}$ OPC and OL (+T3-DIV4) lysate (0 - 8 μg; x-axis; STAR Methods). (B) Normalized β-glucuronidase activity (Units/mg Lysate) for OPC = 8.71 U / mg ± 0.27 and OL = 7.97 U / mg ± 0.18; t-test: $t_{(2)}$ = 2.18; p = 0.16). (C) β-glucuronidase activity in multiple brain regions (cerebral cortex, corpus callosum, striatum and cerebellum) of P21 THAP1 N-CKO (*Thap1*$^{flx/-}$; nestin-*Cre*$^{+}$) and control (*Thap1*$^{+/flx}$) mice. Bar graph shows mean ± SEM values of normalized β-glucuronidase activity Units/Lysate (mg). Cerebral cortex - *Thap1*$^{+/flx}$ = 10.32 U / mg ± 0.43 ; N-CKO = 4.76 U / mg ± 0.106; t-test: p < 0.0001; corpus callosum - *Thap1*$^{+/flx}$ = 13.49 U / mg ±

0.33 ; N-CKO = 5.38 U / mg ± 0.129; t-test: $p < 0.0001$; striatum - *Thap1*$^{+/flx}$ = 9.98 U / mg ± 0.183 ; N-CKO = 5.10 U / mg ± 0.10; t-test: $p < 0.0001$ and cerebellum - *Thap1*$^{+/flx}$ = 14.97 U / mg ± 0.552 ; N-CKO = 5.58 U / mg ± 0.117; t-test: $p < 0.0001$) from P21 mice. (D) Pearson's coefficient value shows ~ 95% colocalization (R = 0.94 ± 0.009; multiple region of interest (ROI) for N=50 cells) for CS-GAG (CS-56) and lysosomes (LAMP1) in OL (DIV4). Graph demonstrating co-localization of CS-56 and LAMP1 pixels for the image represented in FIg. 5N calculated using Image J.

**Figure S5 related figure 5. β-glucuronidase overexpression in CNS tissue.**
β-glucuronidase activity in multiple brain regions (cerebral cortex, corpus callosum, striatum and cerebellum) of P21 *Thap1*$^{+/flx}$ and *Thap1*$^{+/flx}$;Tg$^{GUS}$ mice. Bar graph shows mean ± SEM values of normalized β-glucuronidase activity Units/Lysate (mg). Cerebral cortex - *Thap1*$^{+/flx}$ = 10.32 U / mg ± 0.43 ; *Thap1*$^{+/flx}$;Tg$^{GUS}$ = 2402 U / mg ± 120.6; t-test: $p < 0.0001$; corpus callosum - *Thap1*$^{+/flx}$ = 13.49 U / mg ± 0.33 ; *Thap1*$^{+/flx}$;Tg$^{GUS}$ = 2208 U / mg ± 140.2; t-test: $p < 0.0001$; striatum - *Thap1*$^{+/flx}$ = 9.98 U / mg ± 0.183 ; *Thap1*$^{+/flx}$;Tg$^{GUS}$ = 2056 U / mg ± 110.3; t-test: $p < 0.0001$ and cerebellum - *Thap1*$^{+/flx}$ = 14.97 U / mg ± 0.552 ; *Thap1*$^{+/flx}$;Tg$^{GUS}$ = 2078 U / mg ± 724.9; t-test: $p < 0.0001$.
