## Supplemental Table S3 for "THAP1 Modulates Oligodendrocyte Maturation by Regulating ECM Degradation in Lysosomes"

| **Table S3**: THAP1 regulated genes common to ECM metabolism | | | |
|---|---|---|---|
| Input | MGI Gene/ Marker ID | Name | Entrez Gene ID |
| Gaa | MGI:95609 | glucosidase, alpha, acid | 14387 |
| Slc2a1 | MGI:95755 | solute carrier family 2 (facilitated glucose transporter), member 1 | 20525 |
| Slc26a2 | MGI:892977 | solute carrier family 26 (sulfate transporter), member 2 | 13521 |
| St3gal6 | MGI:1888707 | ST3 beta-galactoside alpha-2,3-sialyltransferase 6 | 54613 |
| Agrn | MGI:87961 | agrin | 11603 |
| Glce | MGI:2136405 | glucuronyl C5-epimerase | 93683 |
| Gusb | MGI:95872 | glucuronidase, beta | 110006 |
| Hmmr | MGI:104667 | hyaluronan mediated motility receptor (RHAMM) | 15366 |
| Hexa | MGI:96073 | hexosaminidase A | 15211 |
| Hs6st1 | MGI:1354958 | heparan sulfate 6-O-sulfotransferase 1 | 50785 |
| Hs3st3b1 | MGI:1333853 | heparan sulfate (glucosamine) 3-O-sulfotransferase 3B1 | 54710 |
| Phka1 | MGI:97576 | phosphorylase kinase alpha 1 | 18679 |
| Hs6st2 | MGI:1354959 | heparan sulfate 6-O-sulfotransferase 2 | 50786 |
| Chsy3 | MGI:1926173 | chondroitin sulfate synthase 3 | 78923 |
| Chst5 | MGI:1931825 | carbohydrate (N-acetylglucosamine 6-O) sulfotransferase 5 | 56773 |
| Sord | MGI:98266 | sorbitol dehydrogenase | 20322 |
| Pfkl | MGI:97547 | phosphofructokinase, liver, B-type | 18641 |
| Eno2 | MGI:95394 | enolase 2, gamma neuronal | 13807 |
| Cd63 | MGI:99529 | CD63 antigen | 12512 |
| Fzd8 | MGI:108460 | frizzled class receptor 8 | 14370 |
| Ctnnb1 | MGI:88276 | catenin (cadherin associated protein), beta 1 | 12387 |
| Msn | MGI:97167 | moesin | 17698 |
| Trp53 | MGI:98834 | transformation related protein 53 | 22059 |
| Ptk2 | MGI:95481 | PTK2 protein tyrosine kinase 2 | 14083 |
| Itga2 | MGI:96600 | integrin alpha 2 | 16398 |
| Flna | MGI:95556 | filamin, alpha | 192176 |
| Actb | MGI:87904 | actin, beta | 11461 |
| Erbb3 | MGI:95411 | erb-b2 receptor tyrosine kinase 3 | 13867 |

| | | | |
|---|---|---|---|
| Vav3 | MGI:1888518 | vav 3 oncogene | 57257 |
| Itgb3 | MGI:96612 | integrin beta 3 | 16416 |
| Itpr2 | MGI:99418 | inositol 1,4,5-triphosphate receptor 2 | 16439 |
| Mdm2 | MGI:96952 | transformed mouse 3T3 cell double minute 2 | 17246 |
| Extl3 | MGI:1860765 | exostosin-like glycosyltransferase 3 | 54616 |
| Nrxn1 | MGI:1096391 | neurexin I | 18189 |
| P4hb | MGI:97464 | prolyl 4-hydroxylase, beta polypeptide | 18453 |
| Nid1 | MGI:97342 | nidogen 1 | 18073 |
| Tgfb3 | MGI:98727 | transforming growth factor, beta 3 | 21809 |
| Capn7 | MGI:1338030 | calpain 7 | 12339 |
| Timp2 | MGI:98753 | tissue inhibitor of metalloproteinase 2 | 21858 |
| Col16a1 | MGI:1095396 | collagen, type XVI, alpha 1 | 107581 |
| Sparc | MGI:98373 | secreted acidic cysteine rich glycoprotein | 20692 |
| Pcolce2 | MGI:1923727 | procollagen C-endopeptidase enhancer 2 | 76477 |
