## Supplemental Table S4 for "THAP1 Modulates Oligodendrocyte Maturation by Regulating ECM Degradation in Lysosomes"

| **Table S4**: Primer sequences used for genotyping | | |
|---|---|---|
| Gene | Forward | Reverse |
| *Thap1* | GCATAGGACAGAGCCTTTCAG | GATGCCAATACCTGATTGGAG |
| | TGCTGGGTGTTGGAAAATAA | |
| *Cre* | CTAGGCCACAGAATTGAAAGATCT | GTAGGTGGAAATTCTAGCATCATCC |
| | GCGGTCTGGCAGTAAAAACTATC | GTGAAACAGCATTGCTGTCACTT |
| $Tg^{GUS}$ | CTA GGC CAC AGA ATT GAA AGA TCT | GTA GGT GGA AAT TCT AGC ATC ATC C |
| | CTG TGG CTG TCA CCA AGA GC | GGA CAC TCA TCG ATG ACC AC |
